# MOAHIT: a web tool for visualizing tumor multi-omics data with human anatomy heatmaps

**DOI:** 10.1101/2022.09.07.506938

**Authors:** Chaozheng Zhou, Anqi Lin, Mengyao Li, Chang Qi, Jian Zhang, Peng Luo

**Affiliations:** Department of Oncology, Zhujiang Hospital, Southern Medical University, Guangzhou, Guangdong, China; Key Laboratory of Breast Cancer, Department of Breast Surgery, Fudan University Shanghai Cancer Center, Shanghai, China

## Abstract

Multi-omics data plays an important role in cancer research, helping clinicians to better explore drug targets and biomarkers. At present, there exist several databases including TCGA (the Cancer Genome Atlas) and GDSC (Genomics of Drug Sensitivity in Cancer), which contain multi omics data on multiple cancer species, as well as various web tools for analyzing the multi-omics data, which are widely used in oncology research. Tumor heterogeneity is a widespread phenomenon, reflected by differences in multi-omics data of cancers originating from different tissues and organs. However, there is a lack of convenient analysis and visualization tools to explore tumor heterogeneity. As a result, we developed a web tool called Multi-Omics Anatomy Heatmap in Tumors (MOAHIT) based on shiny programming. This tool enables users to analyze and visualize the heterogeneity of human tissues. In addition, MOAHIT enables users to produce a beautiful human anatomy heatmap to visualize the heterogeneity between distinct cancers in the multi-omics data.

## Introduction

Multi-omics data includes genomic, proteomic, transcriptome, metabonomic, and epigenetic data[1]. Bioinformatics analysis based on multi-omics data can be used to explore complex biological problems, including development, cell response and disease. By linking genotypes with phenotypes, it can also be used to discover drug targets and biomarkers[2]. Multi-omics analysis is a popular approach in tumor research that can help researchers better connect the tumor phenotypes with observations of tumor tissue, supporting the identification of new tumor therapeutic targets and screen related markers[3][4].

Tumor heterogeneity is a common condition, where tumors originating from different sites can show different morphological and surface characteristics such as differing cell morphology, gene expression, cell metabolism, proliferation and metastasis[5]. This heterogeneity can be observed across different cells within a tumor, and also exists between tumors of a different tissue or organ origin. Multi-omics data is a good representation of tumor heterogeneity, reflecting differences between cancers in different tissues and organs[6]. However, there is a lack of an intuitive and convenient analysis and visualization tool for the study of tumor heterogeneity between different tissues and organs.

With the recent extensive development of tumor multi-omics research, many public databases have been created to store the multi-omics data collected from both cancer patients and normal human tissues. Some major examples include the TCGA database (the Cancer Genome Atlas), which stores tumor samples of more than 20000 individuals representing 33 different cancer types[7], the ICGC (International Cancer Genome Consortium) database for large-scale genomic research, which coordinates 76 cancer projects in 21 primary cancer sites from more than 20000 donors[8], the CCLE (Cancer Cell Line Encyclopedia) database, which stores multi-omics data and drug sensitivity data for several cancer cell lines[9], and the GDSC database (Genomics of Drug Sensitivity in Cancer)[10]. In addition, several databases collect tumor multi-omics data, such as METRBIC (Molecular Taxonomy of Breast Cancer International Consortium)[11] and TARGET[12]. Currently, there are many kinds of online web analysis tools or databases available for tumor multi-omics data, which are widely used in cancer research. A few examples of these include UCSC Xena[13], cBioPortal[14], HPA(Human Protein Atlas)[15], Gene Expression

Profiling Interactive Analysis(GEPIA)[16] and GEPIA2[17]. These web tools can be used for multi-omics data analysis and visualization based on published tumor multi-omics data, with applications such as differential gene expression analysis, gene set enrichment analysis (GSEA) and immune infiltration analysis. However, these tools are not fully suitable for the analysis or visualization of the heterogeneity of cancer multi-omics data in different tissues and organs.

As a result, we have developed a multi-organ visualization tool named Multi-Omics Anatomy Heatmap in Tumor (MOAHIT), which can be used to visualize the heterogeneity between cancers originating from different tissues or organs.

### Methods

MOAHIT web tool is an online analysis web tool based on R language and Shiny programming[18]. It is completely free and does not require a login or registration. Users can visualize published multi-omics data, such as multi omics data stored from TCGA[7], GTEx[19] or GDSC[10] through MOAHIT, or they can upload their own data to the web and customize the visualization results. The User Interface (UI) of MOAHIT is built based on the DashBoard framework, with the server and interactive data constructed using R. Data tables in MOAHIT are displayed using the DT package, which enables viewing, filtering, and downloading of data. Images are generated using the renderplot function, and users can download PDF files of the visualization results. All the code and visual images are built using R software (version. 4.0). Figure 1 shows the architecture flow chart of MOAHIT.

**Figure 1:**
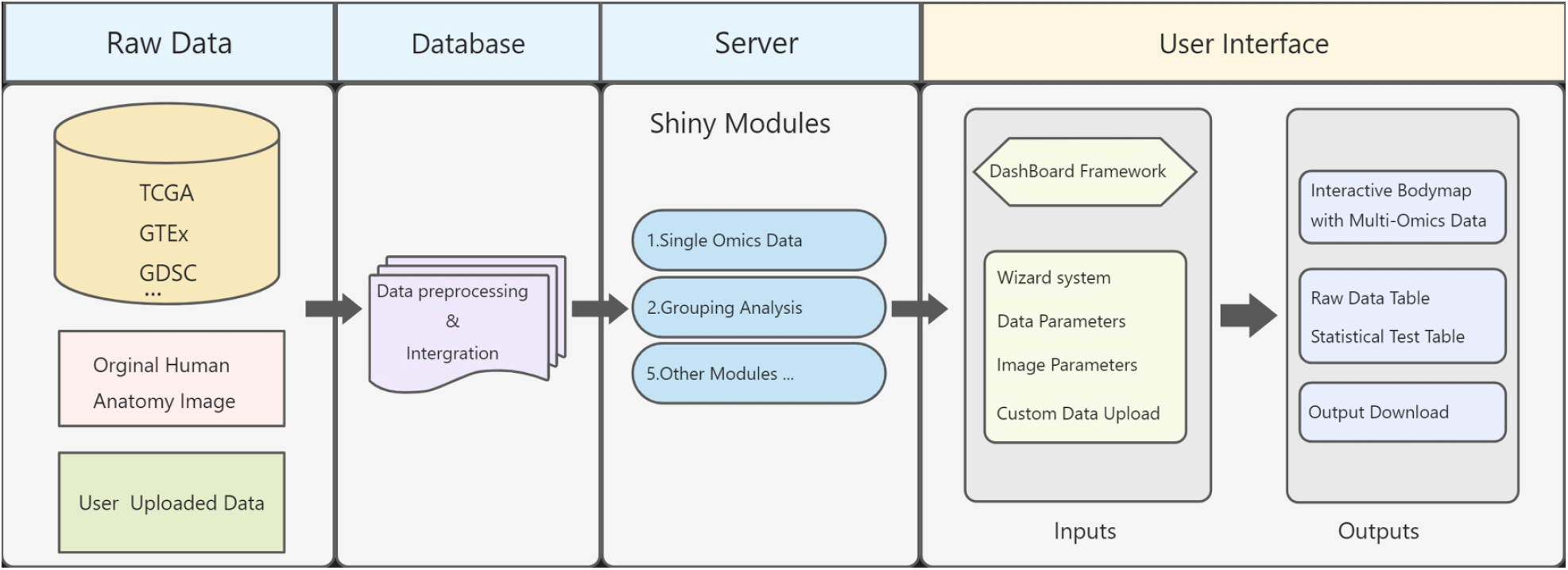
The framework and technical route of MOAHIT.

For published oncology multi-omics data, we collect and download data from the GTEXx database [19] and TCGA Pan-Cancer data [20]. The data included gene expression data, DNA methylation data, copy number variation data, genomic data, and proteome data. All data was stored in the form R.data and stored on our own server.

The human anatomy map in MOAHIT is an original creation by our team, which provides the image basis for MOAHIT to provide exquisite human anatomy heatmap with multi omics data. Its function mainly includes three parts: Single Omics Data, Comparison Analysis, and Custom Analysis. Each part can achieve unique functions and provide users with more analysis options to better meet the requirements of analyzing and visualizing tumor heterogeneity.

MOAHIT version beta 1.0 is now successfully deployed on our server for free use by researchers (https://smuonco.shinyapps.io/MOAHIT/).

## Results

### Welcome

MOAHIT includes a simple graphical interface to introduce the usage of each part of the function, so that users can quickly learn how to use the web tool. MOAHIT currently provides two basic function modules and their functionality is described in the welcome interface. Module 1 is used to visualize data provided by the user himself, named “Visualize your own data”. Module 2 is used to visualize pan-cancer multi-omics data, named “Pancancer multi-omics”. In the bottom of the welcome interface provides version update information, as well as contact information for the development team. This makes it easier for users to understand what changes have taken place in each version and how to give feedback or suggestions to the development team (Figure 2).

**Figure 2:**
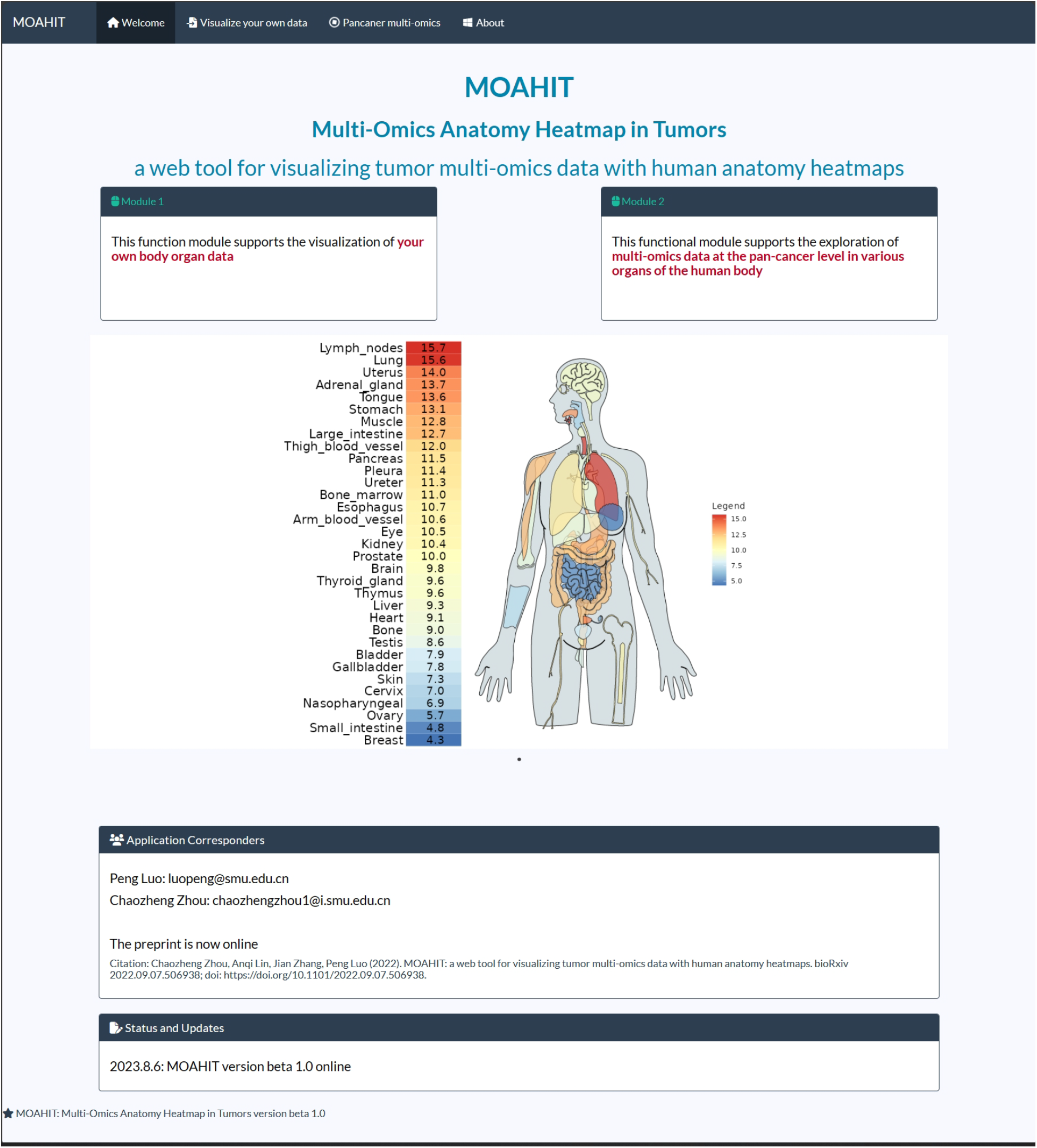
Welcome interface with version information and contact information in the lower left corner.

### Module 1: visualize your own data

#### Custom Analysis

In this module, users can organize their own data into a table (txt or csv format only) containing the specific organ name along with the expression values of the corresponding multi-omics data. We have provided three styles of example data which have different types of variable. Besides, the example data is displayed in the upper left of the interface and is editable for visualizing. Next, the user can click the “Browse” button to upload the data to the server. After previewing the data, users can then click the “Plot” button to visualize their data. It is also possible for the user to modify the visualization details in the “Advanced Plot Parameter” column, enabling them to alter features such as color palette, plot title, legend title and anatomy outline color. Once analysis and visualization are complete, the user can download the image by clicking the “Download Plot” button (Figure 5). Details about parameters in the “Upload and Plot” and “Advanced Plot Parameter” columns can be reached by clicking the question mark icon (Figure 3).

**Figure 3:**
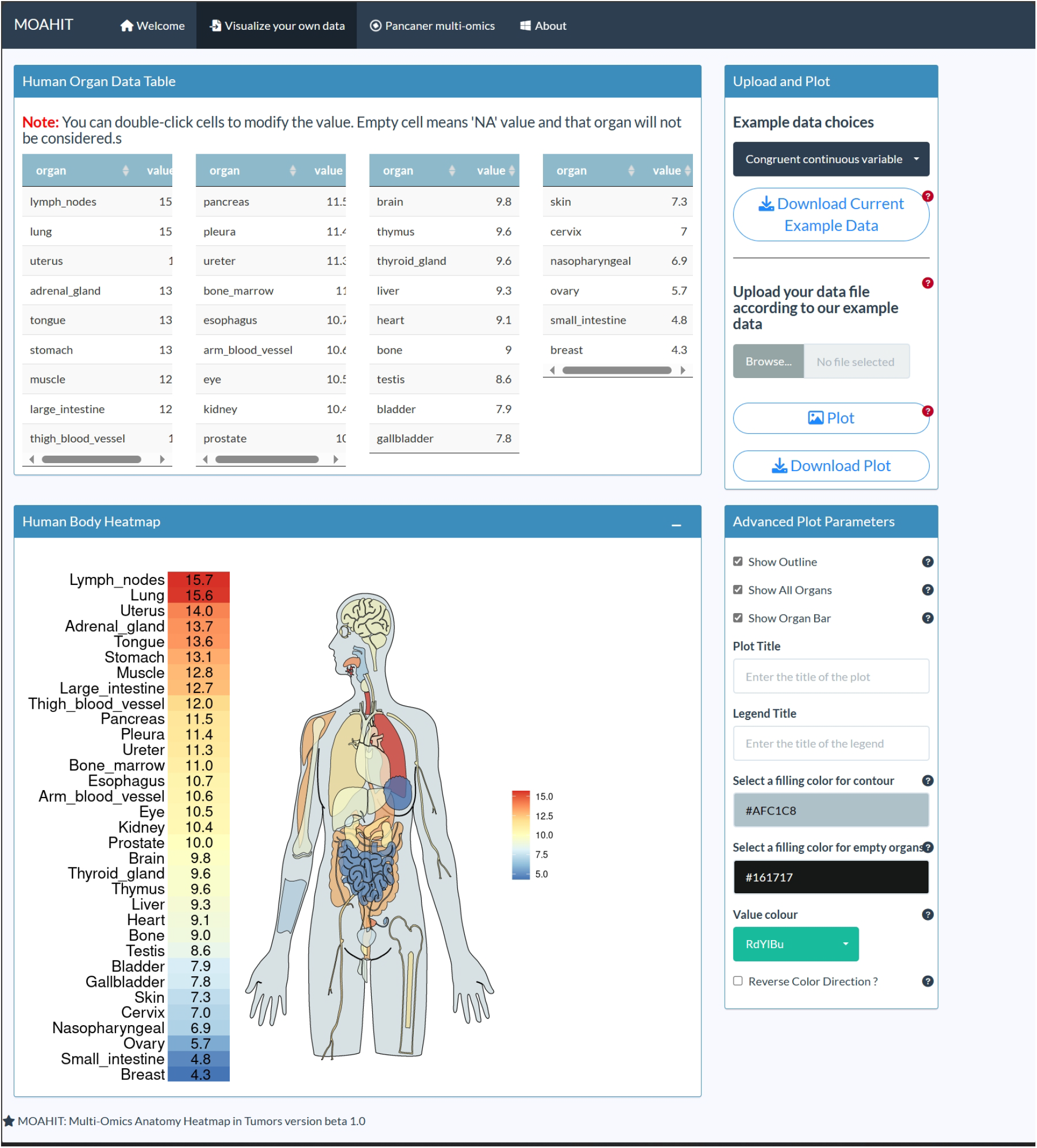
Functional display of the module 1 of MOAHIT named “visualize your own data”. In module 1, users can upload and visualize their data through MOAHIT. Click “Browse” button in the upper right corner to upload your custom data in csv or text format. After uploading, the uploaded data can be previewed in the data table in the upper left corner. By clicking the “Plot” button, the human organ anatomy heat map can be obtained, and the visualization results can then be modified through the “Advanced Plot Parameters” column. Finally, users can download the result by clicking “Download Plot” button.

In this module, users can visualize various types of data (including continuous variables, categorical variable types, etc.) for up to 33 human organs, and can modify the features of the human anatomy heatmap with a high degree of flexibility.

### Module 2: pancancer multi-omics data analysis

#### Single Omics Data

In the “Basic Analysis” interface of MOAHIT, users first need to select the type of multi-omics data to be displayed and the organ, then select the target gene or protein to determine the target molecule to be studied. By clicking the analysis button, users can obtain the single-omics expression data of the target gene in the selected cancer type, as well as an anatomical heat map of different human organs. This analysis can visualize multiple types of multi-omics data from the TCGA database including gene expression, gene mutation frequency, gene copy number variation, and gene methylation and protein expression, with 16 organs corresponding to 33 cancer species available for visualization. Users can then further customize the visualization results, changing the title, color and other image parameters. Once satisfied with the result, users can then download the data corresponding to the selected content as well as the visualization results by clicking on the “Download” button (Figure 4A).

**Figure 4:**
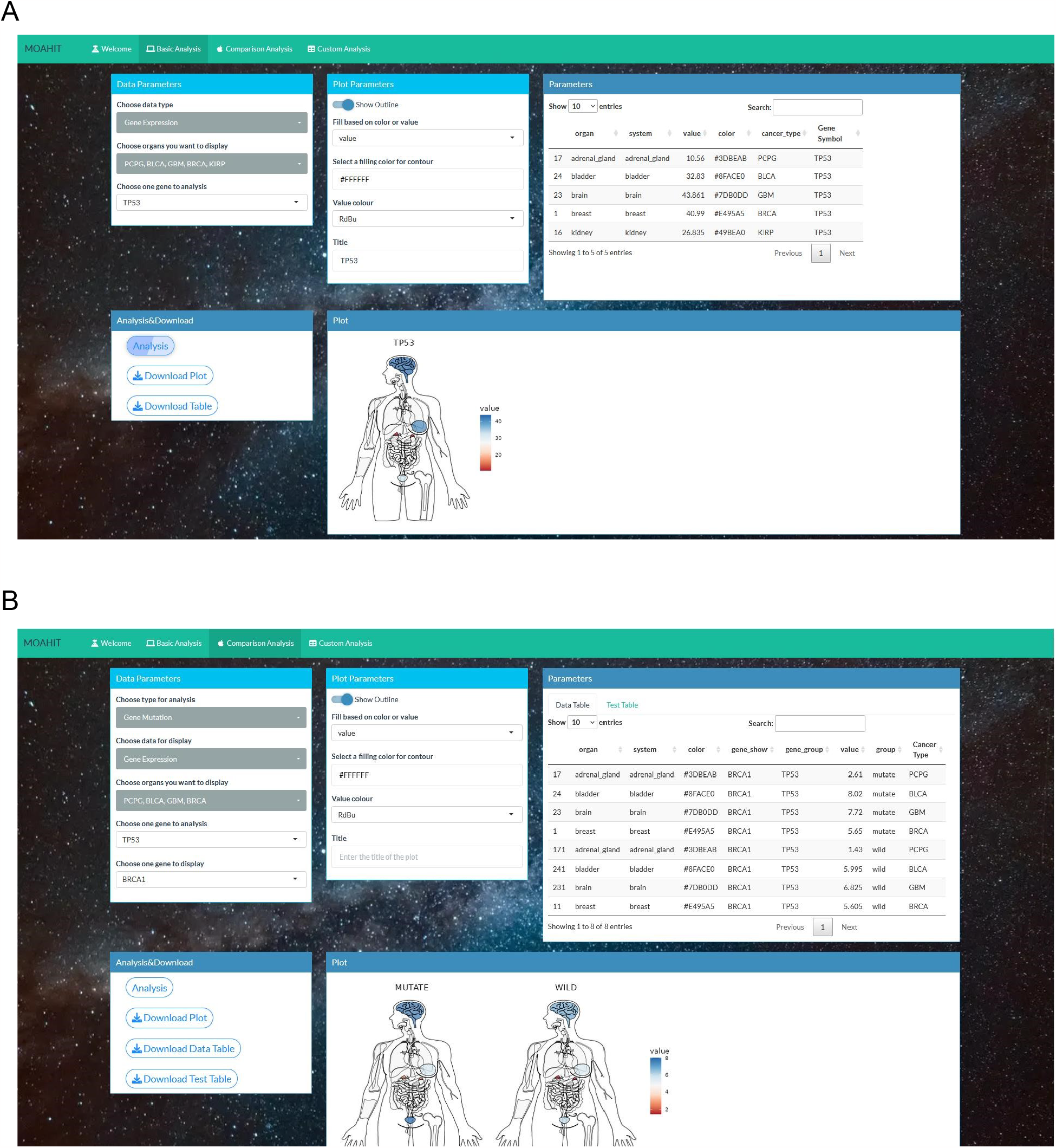
Functional display of the module 2 of MOAHIT named “pancancer multi-omics data analysis”. (A) Basic Analysis module. In the example, the expression data in the data parameters box is first selected, then the cancer types to be displayed as PCPG (Pheochromocytoma and Paraganglioma) representing adrenal gland, BLAC (Bladder Urothelial Carcinoma) representing bladder, BGM (Glioblastoma Multiforme) representing brain, BRCA (Breast Invasive Carcinoma) representing breast, and KIRP (Kidney Renal Papillary Cell Carcinoma) representing kidney. Finally, the target gene TP53 is selected, and the Analysis button can be clicked to display the visualized results in the plot box, showing the expression level of the TP53 gene in the organs represented by the four cancer species. Users can modify the visualization results in the plot parameter box. A data frame showing the selected data is visible on the top right. (B) Functional display of Comparison Analysis module. In the example, the samples were divided into mutation group and wild group according to the mutation status of TP53 gene. The cancer types to be displayed are PCPG representing adrenal gland, BLCA representing bladder, BGM representing brain, BRCA representing breast, and KIRP representing kidney. After clicking the Analysis button, the visualized results can be seen in the plot box, showing the expression level of the BRCA1 gene in the organs represented by the four cancer species with different TP53 mutation status. The statistical test results of the differences between the two groups of data are shown in the top right side of the test table.

#### Comparison Analysis

In the “Comparison Analysis” interface of MOAHIT, users first select the desired target omics type from the multi-omics data for grouping. Using gene mutation data, target genes are selected, and samples are then divided into mutation type (MT) and wild type (WT); alternatively, target molecules can be selected, with the samples then divided into high expression and low expression based on the median expression value of the molecules. Once the grouping results have been obtained, the user can select the data type and target molecule to be displayed. Finally, the user chooses the category of tissues or organs to be displayed. After clicking the “Analysis” button, human organ anatomy heatmaps of the single-omics data of the selected gene are produced based on the grouping result (MT vs WT or high expressed vs low expressed) (Figure 4B).

## Discussion

MOAHIT is an interactive web tool that enables the integration and analysis of tumor multi-omics data and its visualization in human organs, addressing a research need to more easily analyze the heterogeneity between tumors of different origins. Via the “Basic Analysis” module of MOAHIT, researchers can easily explore and visualize the heterogeneity presented in a certain omics data from different tumor types, obtaining a beautiful interactive human anatomy heatmap. Using the “Comparison Analysis” module, samples can be divided into groups according to gene expression or mutation status. It can not only conveniently compare multi-omics expression data in different human tissues and organs in a specific group vertically but can also compare the differences in multi-omics data between two distinct groups horizontally. The “Custom Analysis” module of MOAHIT can solve the research requirement of visualizing the heterogeneity of multi-omics data in tumors originating from different tissues and organs in Pan-cancer analysis. The user image modification function also enables high interactivity with the data and diversity in produced results. As well as its applications in Pan-cancer analysis, this function can also be extended to any situation requiring the mapping of expression values onto human organs. For example, the function could be used to display the number of samples of advanced cancer metastasized to each organ, or to sample the distribution of organs or systems affected by drug-induced adverse events. These functions can enable researchers to explore the heterogeneity between tumors from different tissues and organs using public databases, as well as exploring their own data for Pan-cancer research. It should be noted that MOAHIT can also be used in other scenarios for the visualization of human organ anatomy. A key innovation of this tool, which is currently lacking in other web tools, is the ability of these functions to analyze heterogeneity between tumors from different tissues and organs.

In addition to providing functional advantages in the study of tumor heterogeneity, the human anatomy visualization provided by MOAHIT is an original development of our team, producing simple and beautiful figures. The details of the visualization results can be modified in each functional module to meet the aesthetic needs of researchers and produce figures suitable for publication. The anatomy plot of the human body produced by MOAHIT is similar to the “interactive bodymap” of GEPIA2[17] but is more conveniently organized, provides more visual choices for organ visualization, and is highly customizable.

The current version of the MOAHIT web tool can meet the basic multi-omics data visualization requirement to produce a human organ anatomy map. However, as a newly developed web tool, there is still a lot to improve. We welcome users to contact us to provide any suggestions that enable us to improve this web tool.

## Data availability

R studio is a language and environment for statistical computing available in https://www.r-project.org/. Shiny is an open source collaborative initiative available in the GitHub repository (https://github.com/rstudio/shiny). MOAHIT version beta 1.0 is now successfully deployed on our server for free use by researchers (https://smuonco.shinyapps.io/MOAHIT/).

## Acknowledgements

Special thanks to the English language polishing contributions from TopScience Editing.

## Ethics approval and consent to participate

Not applicable.

## References

[1] I. Subramanian, S. Verma, S. Kumar, A. Jere, and K. Anamika, “Multi-omics Data Integration, Interpretation, and Its Application,” Bioinform. Biol. Insights, vol. 14, 2020, doi: 10.1177/1177932219899051.

[2] B. Vitrinel et al., “Exploiting Interdata Relationships in Next-generation Proteomics Analysis,” Mol. Cell. Proteomics, vol. 18, no. 8 suppl 1, pp. S5–S14, 2019, doi: 10.1074/MCP.MR118.001246.

[3] N. Mouchtouris, R. D. Smit, K. Piper, G. Prashant, J. J. Evans, and M. Karsy, “A review of multiomics platforms in pituitary adenoma pathogenesis,” Front. Biosci. (Landmark Ed., vol. 27, no. 3, Mar. 2022, doi: 10.31083/J.FBL2703077.

[4] G. Nicora, F. Vitali, A. Dagliati, N. Geifman, and R. Bellazzi, “Integrated Multi-Omics Analyses in Oncology: A Review of Machine Learning Methods and Tools.,” Front. Oncol., vol. 10, p. 1030, Jun. 2020 doi: 10.3389/fonc.2020.01030.

[5] A. Marusyk and K. Polyak, “Tumor heterogeneity: causes and consequences,” Biochim. Biophys. Acta, vol. 1805, no. 1, pp. 105–117, Jan. 2010, doi: 10.1016/J.BBCAN.2009.11.002.

[6] S. Turajlic, A. Sottoriva, T. Graham, and C. Swanton, “Resolving genetic heterogeneity in cancer,” Nat. Rev. Genet., vol. 20, no. 7, pp. 404–416, Jul. 2019, doi: 10.1038/S41576-019-0114-6.

[7] J. N. Weinstein et al., “The Cancer Genome Atlas Pan-Cancer analysis project,” Nat. Genet., vol. 45, no. 10, pp. 1113–1120, Oct. 2013, doi: 10.1038/NG.2764.

[8] J. Zhang et al., “The International Cancer Genome Consortium Data Portal,” Nat. Biotechnol., vol. 37, no. 4, pp. 367–369, Apr. 2019, doi: 10.1038/S41587-019-0055-9.

[9] J. Barretina et al., “The Cancer Cell Line Encyclopedia enables predictive modelling of anticancer drug sensitivity,” Nature, vol. 483, no. 7391, pp. 603–607, Mar. 2012, doi: 10.1038/NATURE11003.

[10] W. Yang et al., “Genomics of Drug Sensitivity in Cancer (GDSC): a resource for therapeutic biomarker discovery in cancer cells,” Nucleic Acids Res., vol. 41, no. D1, pp. D955–D961, Nov. 2012, doi: 10.1093/nar/gks1111.

[11] C. Curtis et al., “The genomic and transcriptomic architecture of 2,000 breast tumours reveals novel subgroups,” Nature, vol. 486, no. 7403, pp. 346–352, Jun. 2012, doi: 10.1038/NATURE10983.

[12] X. Ma et al., “Pan-cancer genome and transcriptome analyses of 1,699 paediatric leukaemias and solid tumours,” Nature, vol. 555, no. 7696, pp. 371–376, Mar. 2018, doi: 10.1038/NATURE25795.

[13] M. J. Goldman et al., “Visualizing and interpreting cancer genomics data via the Xena platform,” Nat. Biotechnol., vol. 38, no. 6, pp. 675–678, Jun. 2020, doi: 10.1038/s41587-020-0546-8.

[14] J. Gao et al., “Integrative analysis of complex cancer genomics and clinical profiles using the cBioPortal,” Sci. Signal., 2013, doi: 10.1126/scisignal.2004088.

[15] M. Uhlén et al., “Proteomics. Tissue-based map of the human proteome,” Science, vol. 347, no. 6220, Jan. 2015, doi: 10.1126/SCIENCE.1260419.

[16] Z. Tang, C. Li, B. Kang, G. Gao, C. Li, and Z. Zhang, “GEPIA: a web server for cancer and normal gene expression profiling and interactive analyses,” Nucleic Acids Res., vol. 45, no. W1, pp. W98–W102, Jul. 2017, doi: 10.1093/NAR/GKX247.

[17] Z. Tang, B. Kang, C. Li, T. Chen, and Z. Zhang, “GEPIA2: an enhanced web server for large-scale expression profiling and interactive analysis,” Nucleic Acids Res., vol. 47, no. W1, pp. W556–W560, Jul. 2019, doi: 10.1093/NAR/GKZ430.

[18] J. Doi, G. Potter, J. Wong, I. Alcaraz, and P. Chi, “Web Application Teaching Tools for Statistics Using R and Shiny,” Technol. Innov. Stat. Educ., vol. 9, no. 1, 2016, doi: 10.5070/t591027492.

[19] J. Lonsdale et al., “The Genotype-Tissue Expression (GTEx) project,” Nat. Genet., vol. 45, no. 6, pp. 580–585, Jun. 2013, doi: 10.1038/NG.2653.

[20] J. Liu et al., “An Integrated TCGA Pan-Cancer Clinical Data Resource to Drive High-Quality Survival Outcome Analytics,” Cell, vol. 173, no. 2, pp. 400–416.e11, Apr. 2018, doi: 10.1016/J.CELL.2018.02.052.

